# Integrin β3 Targeting Biomaterial Preferentially Promotes Secretion of bFGF and Proliferation of iPSC-Derived Vascular Smooth Muscle Cells

**DOI:** 10.1101/2021.01.06.425573

**Authors:** Biraja C. Dash, Kaiti Duan, Themis R. Kyriakides, Henry C. Hsia

## Abstract

Human-induced pluripotent stem cell-derived-vascular smooth muscle cells (hiPSC-VSMC) have been shown to promote angiogenesis and wound healing. However, there is a paucity of research on how the extracellular matrix (ECM) microenvironment may impact the hiPSC-VSMC’s function. In this study, our objective was to understand the effect of specific ECM ligand-integrin interaction on hiPSC-VSMC’s paracrine secretion, cell proliferation, and morphology. We here showed a precise modulation of hiPSC-VSMC in a fibronectin functionalized fibrillar collagen scaffold by targeting their integrin β3. The secretion of proangiogenic growth factor, basic fibroblast growth factor (bFGF) was found to be fibronectin dependent via αvβ3 integrin interactions. Also, our data indicate the possible role of a positive feedback loop between integrin β3, bFGF, and matrix metalloproteinase-2 in regulating hiPSC-VSMC’s morphology and cell proliferation. Finally, the secretome with improved proangiogenic activity shows potential for future regenerative applications.

## Introduction

Induced pluripotent stem cells (iPSC) with less immunogenic and ethical concerns are an incredible source for various cardiovascular cells [1, 2]. These iPSC-derived cells mimic the phenotype and function of their in vivo counterparts and carry genetic information from patient. These promising attributes of iPSC technology have now been used to study disease and develop personalized medicine for various cardiovascular diseases [3, 4]. One of the major cardiovascular cells that have been explored recently is vascular smooth muscle cells (VSMC) [5]. VSMC supports the formation of blood vessels and is involved in the pathophysiology of various vascular diseases [6]. Since the last decade, several advancements have been made to generate a large scale of VSMCs of the pure population with very high efficiency [7]. In one such effort, Patsch et al developed a protocol generating more than 80% pure population of VSMC in mere six days [8]. Furthermore, a groundbreaking method developed by Sinha and his colleagues showed differentiation of human iPSC (hiPSC) into developmental origin-specific SMC subtypes form neuroectoderm, lateral plate mesoderm, and paraxial mesoderm using a chemically defined protocol [9]. Similarly, we developed a protocol that can yield millions of hiPSC-VSMC in a matter of 2 weeks. Our protocol does not require a purification step while still leads to the production of more than 95% pure population of VSMC [10]. We further demonstrated that these cells are functionally similar to that of VSMC from lateral plate mesoderm (LM) embryonic origin [10].

The hiPSC-VSMC has shown promising results in the field of regenerative medicine and disease modeling [1, 5, 7]. Human iPSC-VSMC has been used in bioreactors and different scaffold systems to engineering larger blood vessels [11]. A recent report, by Luo et al, demonstrated the production of mechanically robust blood vessels [12]. We and others used hiPSC-VSMC derived from supravalvular aortic stenosis patients to study disease pathophysiology [10, 13, 14]. An elegant study by Misra et al. linked the lack of elastin production and hyperproliferation of SVAS-VSMC to overexpression of integrin β3 [14]. Furthermore, another interesting report demonstrated that LM-derived hiPSC-VSMC is more proangiogenic compared to other subtypes of VSMC and set the foundation for our work on using these hiPSC-VSMC for wound healing [15]. We were the first to demonstrate the secretory function of this hiPSC-VSMC and the abundant number of paracrine factors released from these cells compared to adipose-derived stem cells (ADSC) [16, 17].

ECM based biomaterials have widely been used to modulate stem cell’s secretome [18]. We recently reported that collagen fibrillar density is a potent modulator of paracrine secretion for hiPSC-VSMC [16]. These ECM-based biomaterials initiate a ligand-integrin interaction and switch focal adhesion kinase on, which in turn activates a cascade of signaling that alter stem cell function [19]. Collagen based scaffolds promote different regenerative processes via interactions with α1β1 and α2β1 integrin of cells. Besides collagen, fibronectin is a key ECM-based biomolecule in modulating stem cell function via αvβ3 and *α5β*1 integrin [19]. Fibronectin with both integrin and growth factor-binding motifs in their protein sequence is an attractive biomolecule to engineer integrin-targeting scaffolds [20]. Recent studies have developed complex ECM system utilizing fibronectin and developing methods to present the fibronectin motifs accessible to binding to GF and fibronectins [20–22].

In this study, we sought to determine the effect of fibronectin and integrin β3 interactions in a fibrillar collagen scaffold setting and whether this specific ligand-integrin signaling regulates paracrine secretion, morphology, and proliferation of hiPSC-VSMC (Figure 1). In particular, we examined the secretory profile of important proangiogenic, tissue remodeling, and anti-inflammatory factors produced by hiPSC-VSMC. Such factors may participate in angiogenesis and improve cell adhesion, proliferation, and migration of hiPSC-VSMCs. Furthermore, we investigated the mechanism of fibronectin mediated modulation of iPSC-VSMC. Our study suggests that fibronectin-αvβ3 interaction activates a signaling cascade resulting in upregulation of bFGF, and MMP-2 and a positive feedback loop between integrin β3, bFGF and MMP-2 to promote cell adhesion and proliferation of iPSC-VSMC. The current study suggests a strategy for precise modulation of the hiPSC-VSMC’s secretome to improve cell-mediated regenerative healing.

**Figure 1:**
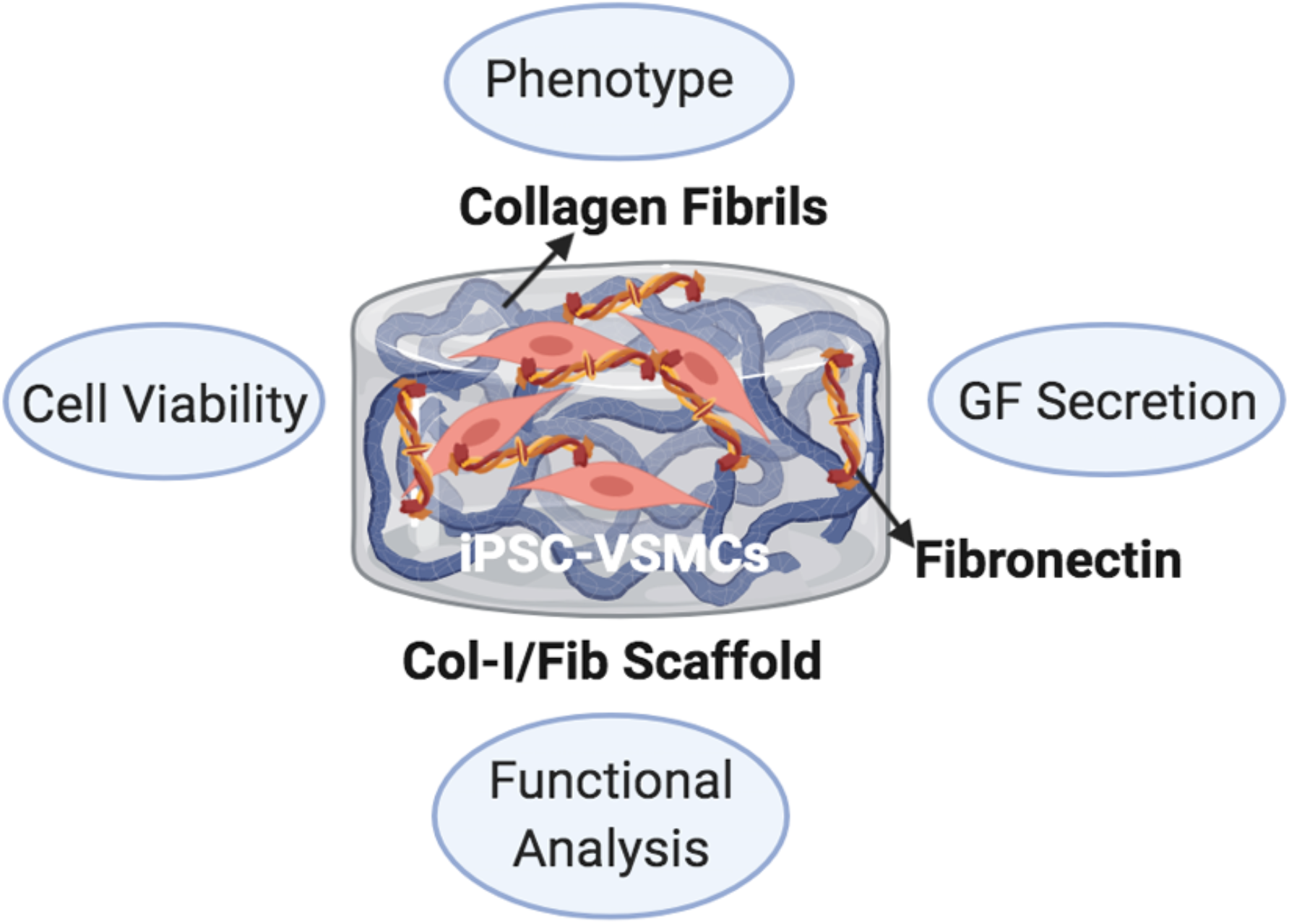
Schematic showing the fabrication and characterization of a Collagen/Fibronectin Scaffold for the modulation of hiPSC-VSMCs proangiogenic factor. Created with BioRender.com

## Methods

### Cell Culture

hiPSC-VSMC generated in a pure population using a previously established protocol were used for all the experiments [10]. The resulting differentiated cells were substantiated using immunofluorescence staining of SM-22α and calponin, SMA, and SM-MHC to check purity. hiPSC-VSMC of passage no more than 7 was used in the experiments. hiPSC-VSMC was grown and prepared on 1% gelatin-coated plates. The medium used smooth muscle growth media (SmGM)-2 with the addition of fetal bovine serum (FBS), with resultant FBS to be 10%. The growth media was replenished every 48 hours, the iPSC-SMC was ready for usage after greater than 80% confluency. Human umbilical endothelial cell (HUVEC) was grown in EGM-2 medium and used after it reached more than 80% confluency.

### Fibronectin collagen scaffold production

The fibronectin functionalized collagen scaffold was constructed in the following steps. Briefly, collagen type-I of the initial concentration of 5mg/ml, fibronectin 1mg/ml, 10xMEM, and 1M NaOH were mixed. The precise ratio of the listed ingredients for each of the formulations can be found in the supplementary table 1. The varying degree of scaffold density was determined and manipulated by the amount of collagen present in the scaffolds. For lower density scaffolds, the rest of the volumes was made out using 1x PBS. For cell encapsulation in the scaffolds, cells were harvested using TrypLE at a concentration of 8000/μl. 200,000 hiPSC-VSMC were then mixed in the neutralized slurry of fibronectin and collagen. The resultant mixture was then distributed as 100ul scaffold aliquots into 96 well cell culture plate. The plate was then incubated for 30 minutes at 37°C for the gelation. 200 μl of Smooth muscle cell media was added on top of the solidified cell scaffolds and subsequently cultured for 3 days. At the end of the 72 hours, the conditioned medium was collected and used for all the downstream processing.

**Table 1:** Various formulations of collagen/fibronectin scaffolds.

| Samples | Density | Collagen | 10x MEM | 1M NaOH | fibronectin<br>1mg/ml | SMC<br>media | SMC<br>8k/ul |
| --- | --- | --- | --- | --- | --- | --- | --- |
| CTRL-1.25mg | 1.25mg | 125ul | 50ul | 2.4ul | 0ul | 300ul | 25ul |
| Fib 25-1.25mg | 1.25mg | 125ul | 50ul | 2.4ul | 25ul | 275ul | 25ul |
| CTRL-2.5mg | 2.5mg | 250ul | 50ul | 3.3ul | 0ul | 175ul | 25ul |
| Fib 25-2.5mg | 2.5mg | 250ul | 50ul | 3.3ul | 25ul | 150ul | 25ul |
| CTRL-4mg | 4mg/ml | 400ul | 50ul | 8.4ul | 0ul | 25ul | 25ul |
| Fib 25-4mg | 4mg/ml | 400ul | 50ul | 8.9ul | 25ul | 0ul | 25ul |
| Fib 50-4mg | 4mg/ml | 400ul | 50ul | 8.9ul | 50ul | 0ul | 25ul |

### Integrin inhibition assay

The hiPSC-VSMC cultures in 2D and 3D scaffolds were incubated with echistatin and ATN-161, potent antagonists for αvβ3 and α5β1 respectively [23, 24], at a concentration of 100 nM concentration for 24 hr on day 2 of culture. The cells and conditioned medium on day 3 were used to perform ELISA and AlamarBlue.

### Immunofluorescence staining

Human iPSC-VSMCs that were cultured on tissue culture plates and scaffolds were verified for phenotype characterization using anti-calponin, anti-SM-22α, anti-SMA, and anti-SMMHC primary antibodies. Selected samples were initially blocked with 5% BSA in 0.25% Triton X for 1 hour at room temperature. Authentication primary antibodies were added for incubation overnight at 4°C. On the next day, the scaffolds were washed three times with PBST (tween20 0.05%), and then the samples were incubated with secondary antibodies tagged with Alexa Fluor^®^ 488 for 2 hours at room temperature. Dapi was used as a counterstain. Z stacks were taken at the interval 1 μm and images were merged for the final image. Finally, the samples were washed in PBST three times at 5 minutes each before fluorescence imaging. The information related to primary and secondary antibodies can be found in tables S1 and S2.

### AlamarBlue assay

hiPSC-VSMC fibronectin induced collagen scaffolds were characterized for viability on day 3 of scaffold methodology as described. First, 200 μl of culturing media was gently aspirated out for future ELISA assays. The scaffolds were then gently washed with 200 μl of 1x PBS. AlamarBlue working solution was made at a 1:10 ratio of AlamarBlue stock to SmGM-2 medium. 100 μl of the AlamarBlue working solution was added to each of the scaffolds and incubated at 37°C for 2 hrs. After the incubation, the plate was read for Fluorescence intensity at 540 nm excitation and 590 emission wavelengths. Relative cell viability was evaluated by dividing them with the collagen scaffold fluorescence intensity value.

### Lactate dehydrogenase assay

Lactate dehydrogenase cytotoxicity assay was performed using an LDH cytotoxicity assay kit (Peirce, Thermofisher) as depicted [16, 25]. Conditioned medium (CM) was used to determine the level of LDH in different samples. Groups that were included in this experiment were: CTRL, fib 25, and fib 50. Absorbance was measured at 490 and 680 nm. To determine LDH activity, the 680nm absorbance value (background signal from the instrument) was subtracted from the 490 nm absorbance. Relative cytotoxicity value to control was determined.

### Live/Dead assay

Cell viability was also assessed via the Live/Dead assay as described. hiPSC-SMC scaffolds were incubated in PBS augmented with 4 μM calcein-AM green (ThermoFisher, USA) and 2 μM ethidium homodimer-1 (ThermoFisher, USA) for 30 minutes. Samples were visualized on an inverted Confocal Laser Scanning Microscope. Z stacks were performed at *1* μm image intervals and images were merged for the final image. Four fields were obtained per scaffold and were imaged for each experiment. Viable and dead cells were counted using ImageJ software. The number of live and dead cells was then approximated.

### Qualitative ELISA

Day 3 culture media collected from the scaffolds were used to perform qualitative ELISA to assess for several pro-angiogenic growth factors VEGF, bFGF, MMP-2, Ang-1, TGF-beta, IL-10 SDF- 1α, and PDGF-AA released from hiPSC-VSMC as described [16, 25]. 2X 90μl of the CM was placed to microwells of 96 well ELISA plates (NUNC MaxiSorp™) as duplicates. The plates were incubated at 4°C overnight. The next day, the plates were washed with PBST (PBS + 0.05% Tween-20) three times and blocked using 5% BSA for 1 hour at room temperature. Then the plates were washed with PBST three times and were incubated with primary antibodies (1:2500) at 4°C overnight. The next day, the plates were washed three times with PBST and incubated with secondary antibodies conjugated with horseradish peroxidase (1:2500) for 2 hours at room temperature, avoid direct lighting whenever possible. The plates were then washed 3 times with PBST before adding 100 μl of TMB substrate solution (Cell Signaling Technology, USA, Catalog: 7004S) to the plates after aspirating out the PBST and was incubated for 25 minutes at room temperature on a plate shaker. 100μl of stop solution (Cell Signaling Technology) was then added to each microwell and absorbance was measured at 450 nm on a plate reader. The information related to primary and secondary antibodies can be found in tables S1 and S2 in the supplementary information. The relative level of growth factor was evaluated by dividing them with the collagen scaffold absorbance value.

### Cell adherence and migration assay

Cell adherence assay for HUVECs was done in a 96 well plate. Briefly, 10, 000 cells were plated on a 96 well plate and cultured for 3 hours in CM supplemented smooth muscle basal medium (SmBM) of a ratio of 1:5. Controls were smooth muscle cell growth medium (SmGM-2), and SmBM. For the migration experiment, trans-well assays were performed using HUVECs as described (Dash et al., 2020). 1 x 10^4^ cells were seeded on the top layer of the trans-wells (8μm pore size trans-wells, Falcon) and incubated for 10 min in cell culture incubators at 37°C. The trans-wells were then placed into 24 well plates and 600μl of respective growth medium with CM were gently added to the wells from the inner side of the trans-well. The trans-wells were then kept in the cell culture incubator for 4 hours. Fixation was done with 100% cold methanol and gently removing the cells from the top layer. Transwells were washed and stained with Dapi. The number of cells migrated to the inner side of the membrane was then quantified by counting the number of Dapi stained nuclei from 4 different fields.

### In vitro network formation assay

The capability of hiPSC-VSMC-CM obtained from in situ scaffolds to induce angiogenesis was evaluated using an in vitro network formation assay as described [25, 26]. HUVECs were cultured until they were 80% confluent in an EGM-2 medium. 100 μl of Matrigel (Corning Life Sciences) was used to evenly coat wells of 96-wells plates. The plates were incubated for 30 min at 37°C to allow the Matrigel to solidify. HUVEC (2×10^5^/ml) that were dislodged in EGM-2 basal medium and 100 μl of this HUVEC containing solution along with 100μl of CM were seeded in each well (n=3). After 6 hours of incubation at 37°C, any floating cells were then removed & the plates were washed twice with PBS and fixed using PFA for brightfield imaging. Images taken from 4 different fields/well were acquired and the number of nodes per field was calculated.

## Results

### Fibronectin mediates proliferation of hiPSC-VSMCs via α_V_β_3_

Human iPSC-VSMC was generated using a previously established protocol [10]. The hiPSC-VSMC was characterized for major smooth muscle cell markers calponin, SM-22α, SM-MHC, and SMA. The cells were highly positive for the aforementioned markers (Figure 2A). The cells were positive for SM-MHC but did not express mature fibers (2A). The hiPSC-VSMC was then seeded on fibronectin-coated plates to determine their proliferation. The cells on days 1, 2, and 3 proliferated on fibronectin and were significantly more compared to the control with no fibronectin (Figure 2B). Most importantly, the hiPSC-VSMC in control wells did not significantly proliferate over three days. Furthermore, the AlamarBlue assay showed a significantly reduced proliferation of hiPSC-VSMCs on fibronectin upon treatment with echistatin, a known antagonist of α_v_β_3_ (Figure 2C).

**Figure 2:**
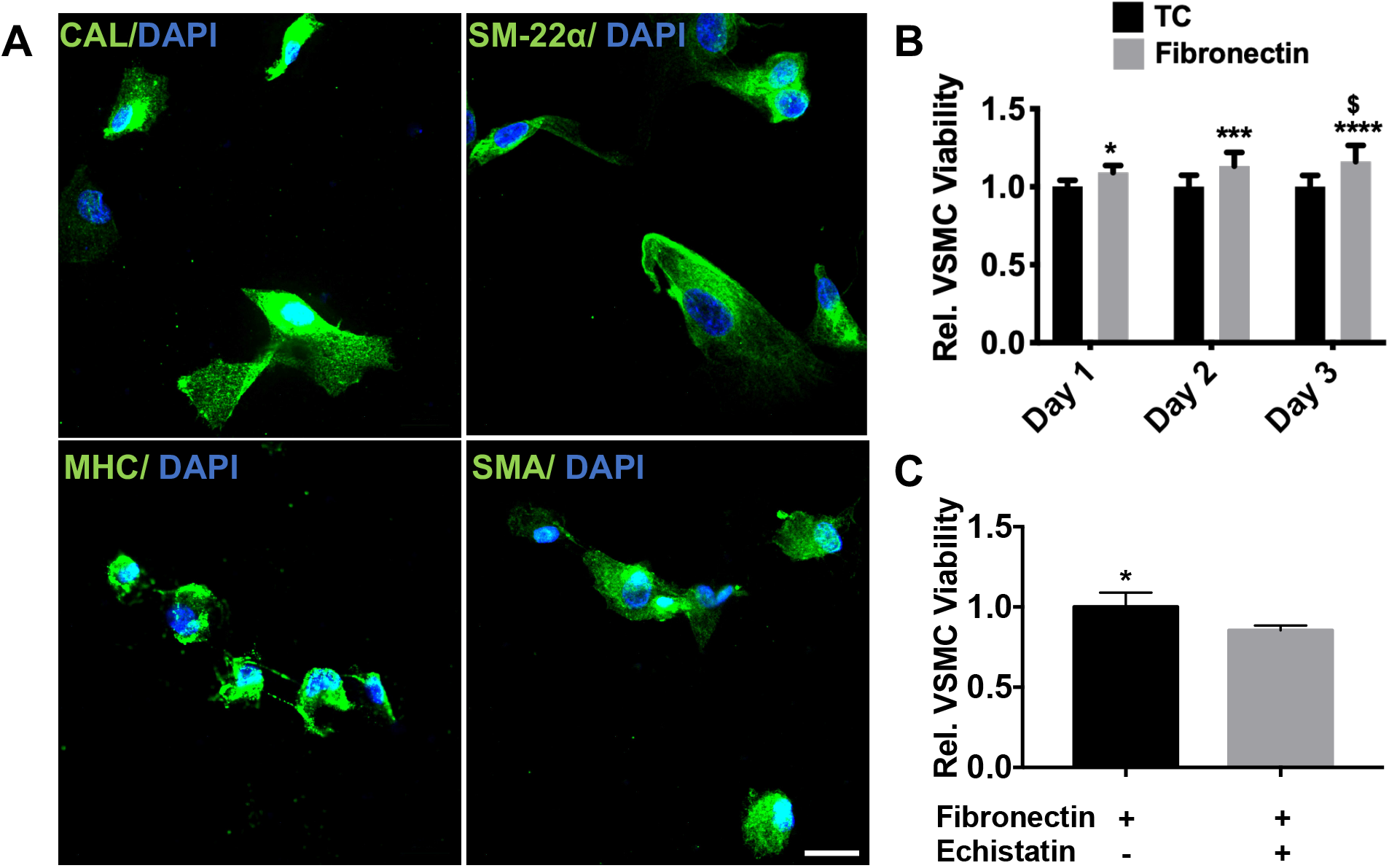
2D characterization of the effect of fibronectin on hiPSC-VSMC’s viability. (A) The hiPSC- VSMCs differentiated using an embryoid body method were characterized using smooth muscle cell markers. Phenotype assessment of hiPSC-VSMCs were performed using immunofluorescence staining with Calponin, SM-22α, smooth muscle myosin heavy chain (SM-MHC) and smooth muscle α actin (SMA) in Green and Dapi stained nuclei in Blue. Scale bar measures 20 μm. AlamarBlue assay showing relative cell viability of hiPSC-VSMCs (B) with and without fibronectin coating for day 1, 2 and 3 and (C) with and without α_v_β_3_ antagonist echistatin (100 nM/mL) on fibronectin coated plates on day 3. * denotes statistical significance differences between the different groups (n=4-12, one-way and two-way ANOVA, *p < 0.05, ***p<0.001, and ****p<0.0001).

The hiPSC-VSMC was then embedded in a 3D collagen scaffold functionalized with fibronectin and was analyzed for cell proliferation, cytotoxicity, and phenotype after three days in culture (Figure 3). The collagen scaffold of 4mg/ml final concentration was used with fibronectin of 50 μg/ml of final concentration. The AlamarBlue assay data showed an enhanced proliferation of hiPSC-VSMC in the fibronectin functionalized scaffolds (Figure 3A). The LDH assay was used to assess the cytotoxicity of the collagen/fibronectin scaffold and control collagen scaffolds. The LDH assay showed a minimal cytotoxic effect of collagen/fibronectin as compared to the control collagen scaffolds (Figure 3B). Furthermore, immunofluorescence staining was performed to characterize the morphology and phenotype of hiPSC-VSMC in the scaffolds (Figure 3C). Immunofluorescence images showed an elongated morphology of hiPSC-VSMC inside collagen/fibronectin scaffolds as compared to collagen control scaffolds. The hiPSC-VSMC maintained their smooth muscle cell phenotype in the hydrogels as shown by staining with calponin and SM-22α, SM-MHC, and SMA (Figure 3C) during three days of in vitro culture. Figure 3D shows immunofluorescence images of Calcein-AM^+^ cells (live) and EthD-1^+^ (dead) cells. The graphs showed an increased number of live hiPSC-VSMC in collagen/fibronectin scaffolds compared to control scaffolds (Figure 3D and E), whereas the ratios of the dead to the total number of cells were found to be similar in both the scaffolds (Figure 3F).

**Figure 3:**
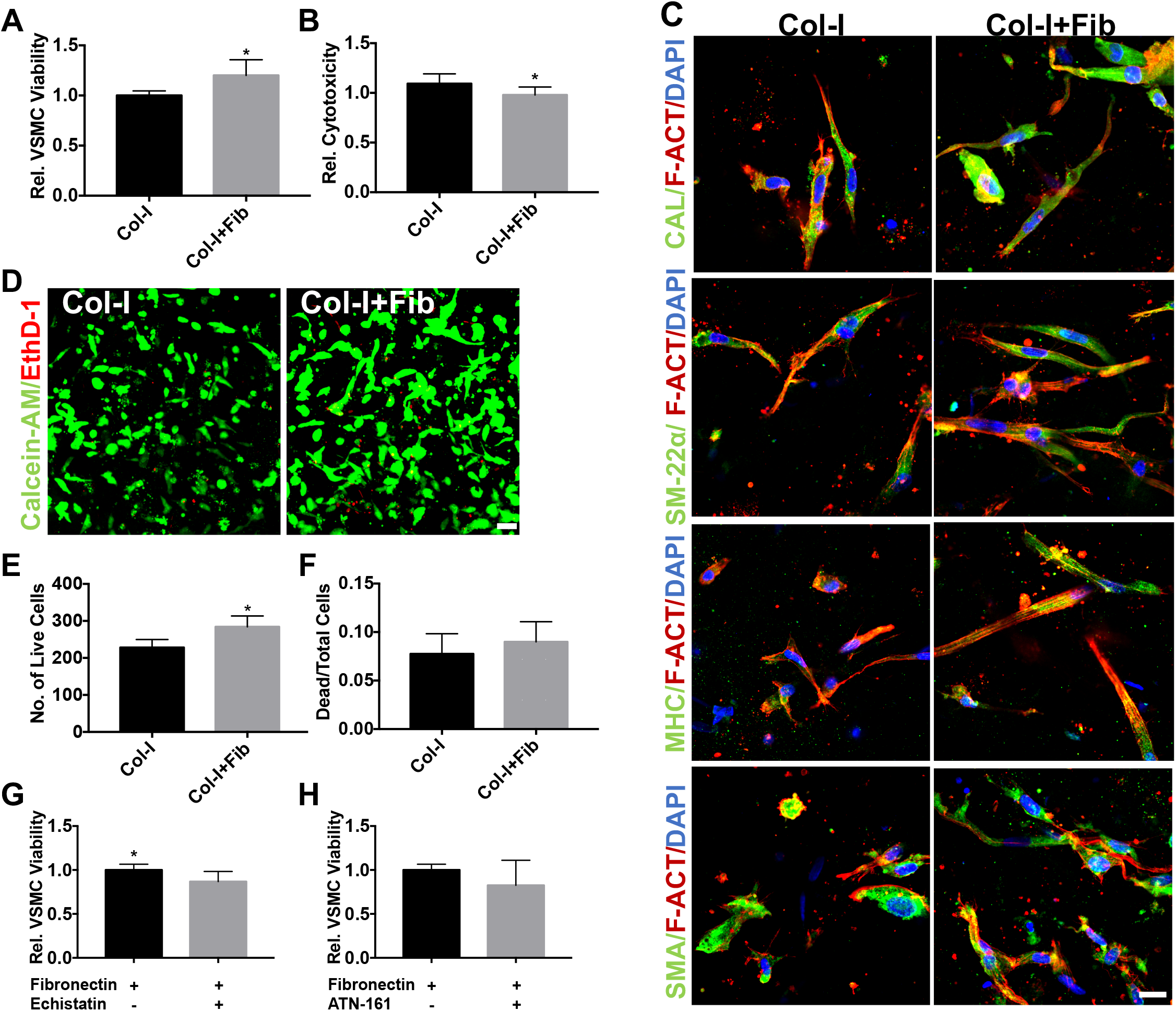
Characterization of hiPSC-VSMC viability and phenotype embedded in the collagen/fibronectin scaffold. (A) AlamarBlue cell viability assay showing relative cell viability of hiPSC-VSMCs embedded in the collagen/fibronectin scaffolds of 4mg/ml of collagen concentration. (B) LDH assay of conditioned medium of the collagen/fibronectin scaffolds. (C) Phenotype assessment of hiPSC-VSMCs embedded in the in situ hydrogels. Immunofluorescence images showing calponin, SM- 22α, smooth muscle myosin heavy chain (SM-MHC) and smooth muscle α actin (SMA) in Green and Dapi stained nuclei in Blue. Scale bar measures 20 μm. (D) Live/Dead assay of the collagen/fibronectin scaffold. Calcein-AM and EthD-1 stains live cells (Green) and dead cells (Red) respectively. Scale bar measures 50 μm. The graph shows (E) the number of Calcein-AM positive live (green) cells and (F) the ratio of dead/total cells. Collagen scaffolds without fibronectin were kept as controls for all the experiments. AlamarBlue assay showing relative cell viability of hiPSC-VSMCs in collagen/fibronectin scaffolds with and without (G) α_v_β_3_ antagonist echistatin and (H) α_5_β_1_ antagonist ATN-161. * denotes statistical significance differences between the different groups (n=3-6, t-test, *p<0.05).

Furthermore, AlamarBlue assay showed a significantly reduced proliferation of hiPSC-VSMC in collagen/fibronectin scaffold in response to echistatin (Figure 3G), whereas we did not observe any effect of ATN-161, a potent inhibitor of α_5_β_1_, on the proliferation of iPSC-VSMC in the collagen/fibronectin scaffold (Figure 3H).

### Fibronectin selectively upregulates bFGF secretion via α_V_β_3_

Qualitative ELISA-based analysis demonstrated the fold change of proangiogenic growth factors VEGF, bFGF, ANG-1, SDF-1α and PDGF-AA, TGFβ, MMP-2, and IL-10 in the CM of scaffolds (Figure 4A). The level of bFGF and MMP-2 was seen to be upregulated in the collagen/fibronectin scaffold. Especially, bFGF was seen to be enhanced 4-5 folds in the case collagen/fibronectin scaffolds compared to the control collagen scaffolds (Figure 4A). The relative level of bFGF was then investigated in collagen/fibronectin scaffolds across different densities of type I collagen. The data showed an enhanced level of bFGF in the higher density (4mg/ml) of type I collagen (Figure 4B). Figure 4C shows an increase in the relative bFGF level with an increase in the amount of fibronectin from 10 to 100 μg/ml with 100 μg/ml releasing the most significant amount of bFGF, while 50ug had a sub-optimal increase in the bFGF. 10 and 25 μg/ml showed no difference and were similar to that of control collagen scaffolds. The treatment with echistatin showed a decrease in the bFGF secretion level compared to no treatment (Figure 4D). However, we did not see any effect of ATN-161 on bFGF release (Figure 4E).

**Figure 4:**
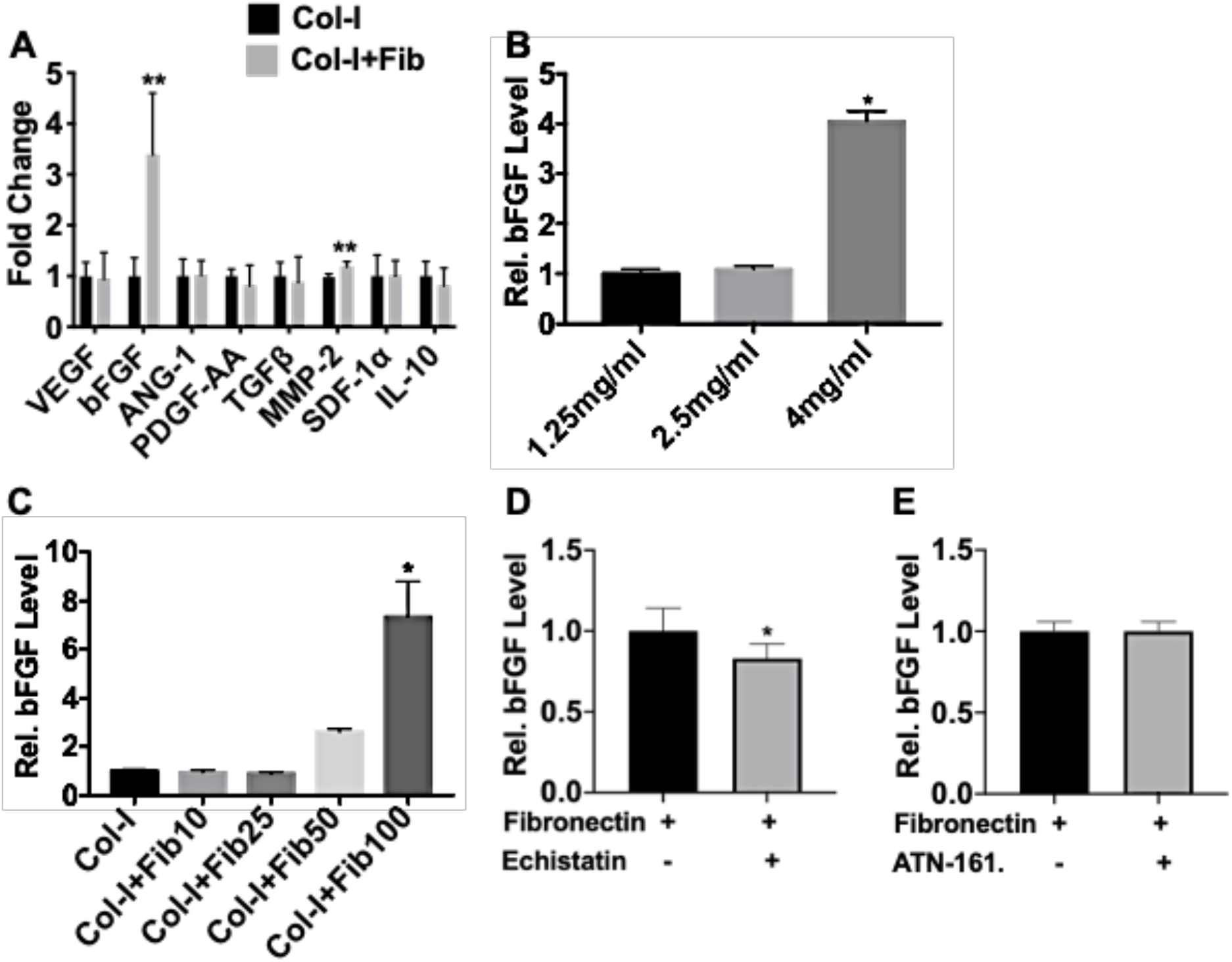
Characterization of pro-angiogenic growth factor secretion from the collagen/fibronectin scaffolds. (A) Qualitative ELISA showing secretion of relative levels of growth factors VEGF, bFGF, ANG-1, PDGF-AA, TGFβ, MMP-2, SDF-1α, and IL-10. ELISA data showing relative bFGF level in (B) response to fibronectin (50 μg/ml) in various densities of collagen scaffold, and (C) different dosages (5, 10, 25, 50 and 100 μg/ml) of fibronectin in a 4mg/ml density of collagen scaffold. Inhibition assay with with and without (D) echistatin and (E) ATN-161 in a 4mg/ml density of collagen scaffold with 100 μg/ml of fibronectin. Collagen scaffolds with and without fibronectin were kept as controls. * denotes statistical significance differences between the different groups (n=3-6, t-test and one-way ANOVA, *p<0.05, **p<0.01).

We saw a comparatively enhanced relative level of MMP-2 secretion in 2.5 mg/ml and 4 mg/ml fibronectin functionalized collagen scaffolds (Figure 5A) compared to control collagen scaffolds. However, we did not see any difference in the MMP-2 secretion between 50 to 100 μg/ml of fibronectin (Figure 5B). Also, incubation with Echistatin and ATN-161 did not show any difference in the MMP-2 secretion in the cells embedded in collagen/fibronectin scaffolds. (Figure 5C and 5D).

**Figure 5:**
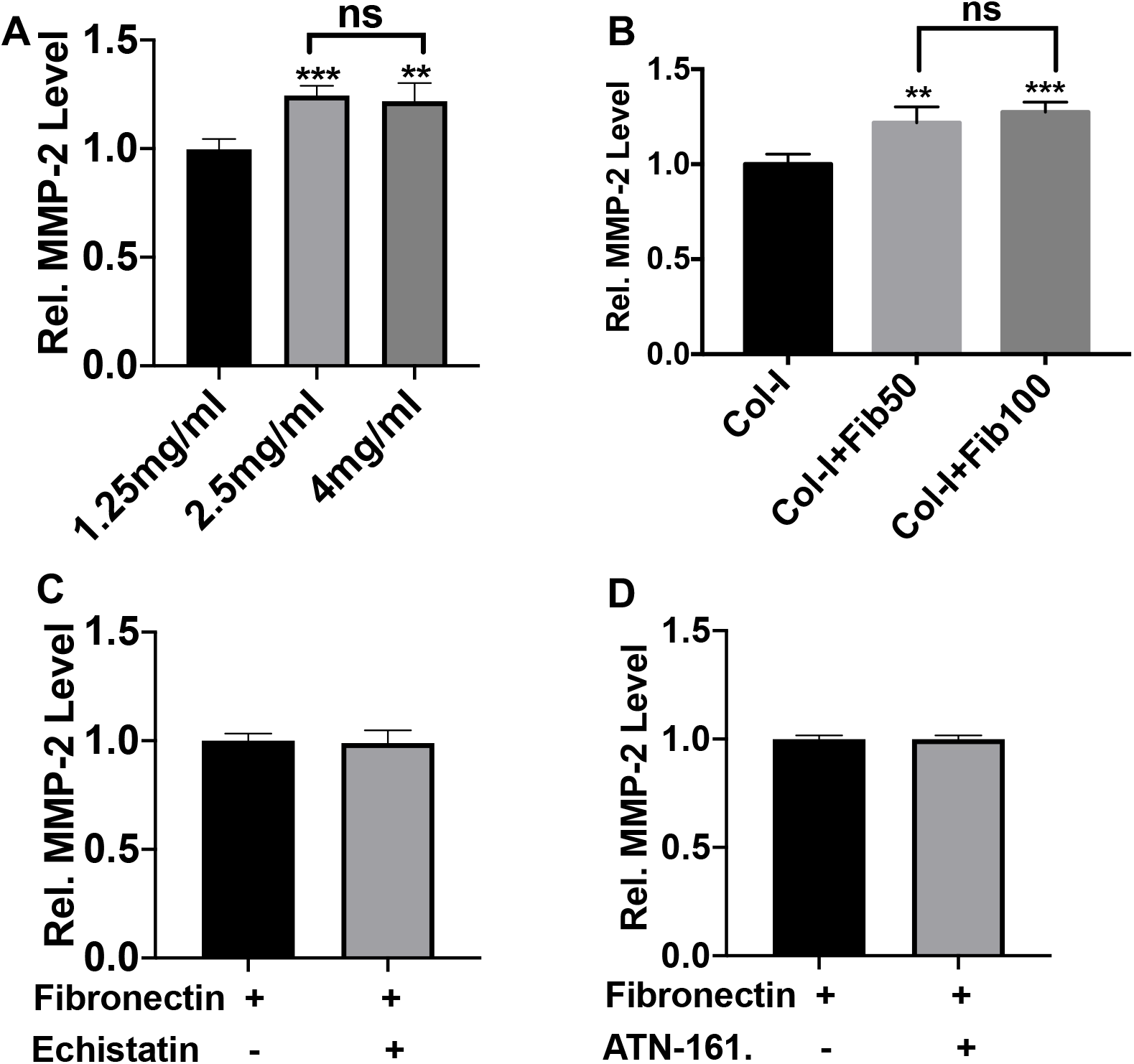
Characterization of MMP-2 secretion. ELISA data showing relative MMP-2 level in response to (A) fibronectin (50 μg/ml) in various densities of 4mg/ml collagen scaffold, (B) different dosages (50 and 100 μg/ml) of fibronectin in a 4mg/ml collagen scaffold, (C) incubation with Echistatin and (D) ATN- 161. Collagen scaffolds with and without fibronectin were kept as controls. * denotes statistical significance differences between the different groups (n=3-6, t-test and one-way ANOVA, **p<0.01, ***p<0.001).

Furthermore, when we cross-linked the collagen/fibronectin scaffold with a PEG-based amine cross-linker, 4S-StarPEG, the release of bFGF was reduced to the level of control scaffolds (Figure 6A). A similar trend was seen for MMP-2 release as well (Figure 6B). There was no difference in the cell viability between cross-linked and non-crosslinked scaffold groups (Figure 6C). The cross-linking of amine groups was investigated and can be seen in figure S1 in the supplementary information. The TNBSA data quantifies free primary amine groups and can be seen lowering in the cross-linked group compared to the control (non-crosslinked).

**Figure 6:**
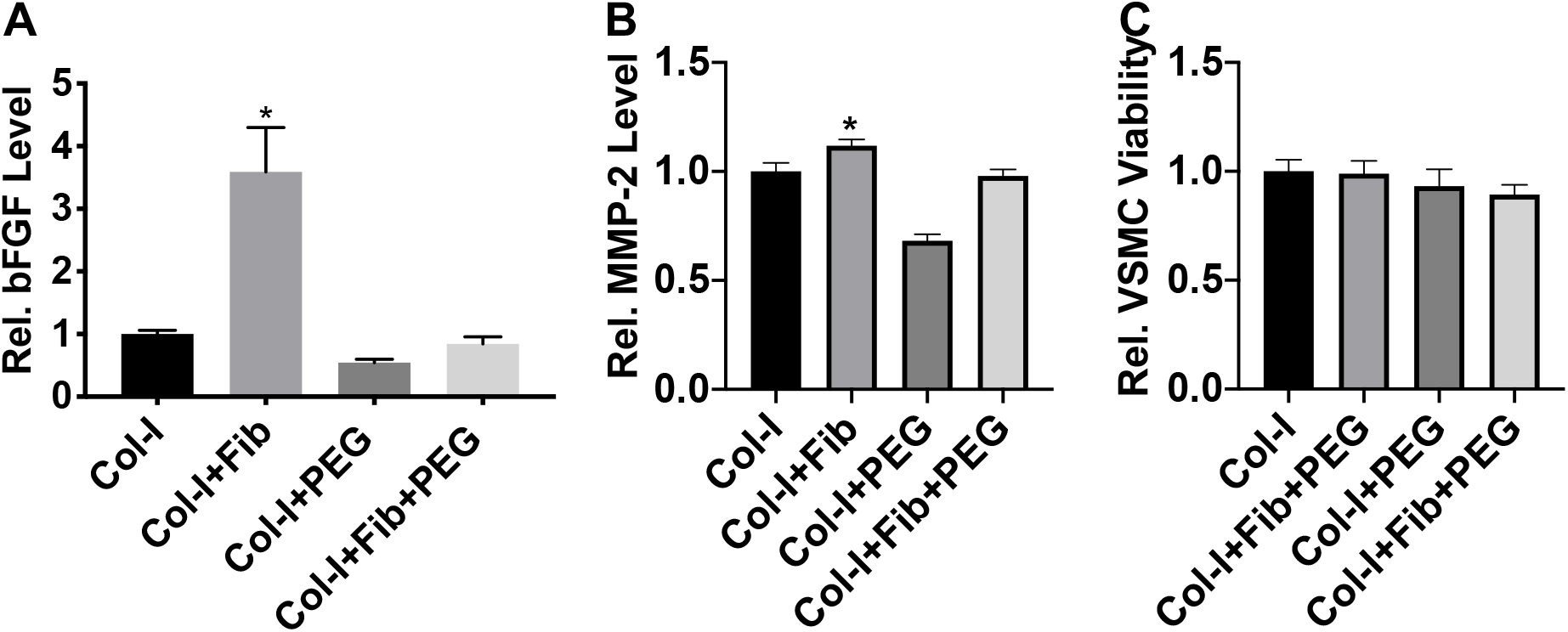
Characterization of the effect of cross-linking on bFGF and MMP-2 secretion. Qualitative ELISA showing secretion of relative level of (A) bFGF and (B) MMP-2 in various formulation of collagen hydrogel: collagen only (Col-I), fibronectin functionalized collagen gel (Col-I+Fib), 4S-StarPEG crosslinked collagen gel (Col-I+PEG) and 4S-StarPEG cross-linked collagen and fibronectin gel (Col- I+Fib+PEG). AlamarBlue assay showing viability of hiPSC-VSMC in aforementioned collagen hydrogel formulations. * denotes statistical significance differences between the different groups (n=4, one-way ANOVA, *p<0.05).

### The conditioned medium from the fibronectin functionalized collagen scaffold promotes endothelial cell adhesion, migration, and network formation

Functional assays of the CM were carried out to determine the bioactivity of the secreted paracrine growth factors from the final collagen/fibronectin scaffolds (Figure 7) and were compared with SmGM-2, positive control endothelial cell growth medium-2 (EGM-2), and negative control endothelial basal medium (EBM). Cell adherence of HUVEC was found to be significantly more in CM compared to control SmGM-2, EGM-2, and EBM medium controls (Figure 7A and B). The HUVECs showed typical cobblestone-like morphology in the case of both CM and EGM-2 after 3 h of seeding (Figure 7A). A similar pattern can be seen in the case of migration of HUVECs towards CM in a transwell migration assay (Figure 7C and D). HUVECs in the case of SmGM-2 and EBM showed reduced migration compared to CM and EGM-2 (Figure 7C and D). And SmGM-2 had enhanced migration compared to the EBM control medium. In vitro network formation assay using HUVECs on Matrigel showed higher nodes/field in the case of both EGM-2 and CM and were significantly more than that of the SmGM-2 and EBM (Figure 7E and F). There was no difference between SmGM-2 and EBM, although an increase can be seen in the case of SmGM-2 it was not significant.

**Figure 7:**
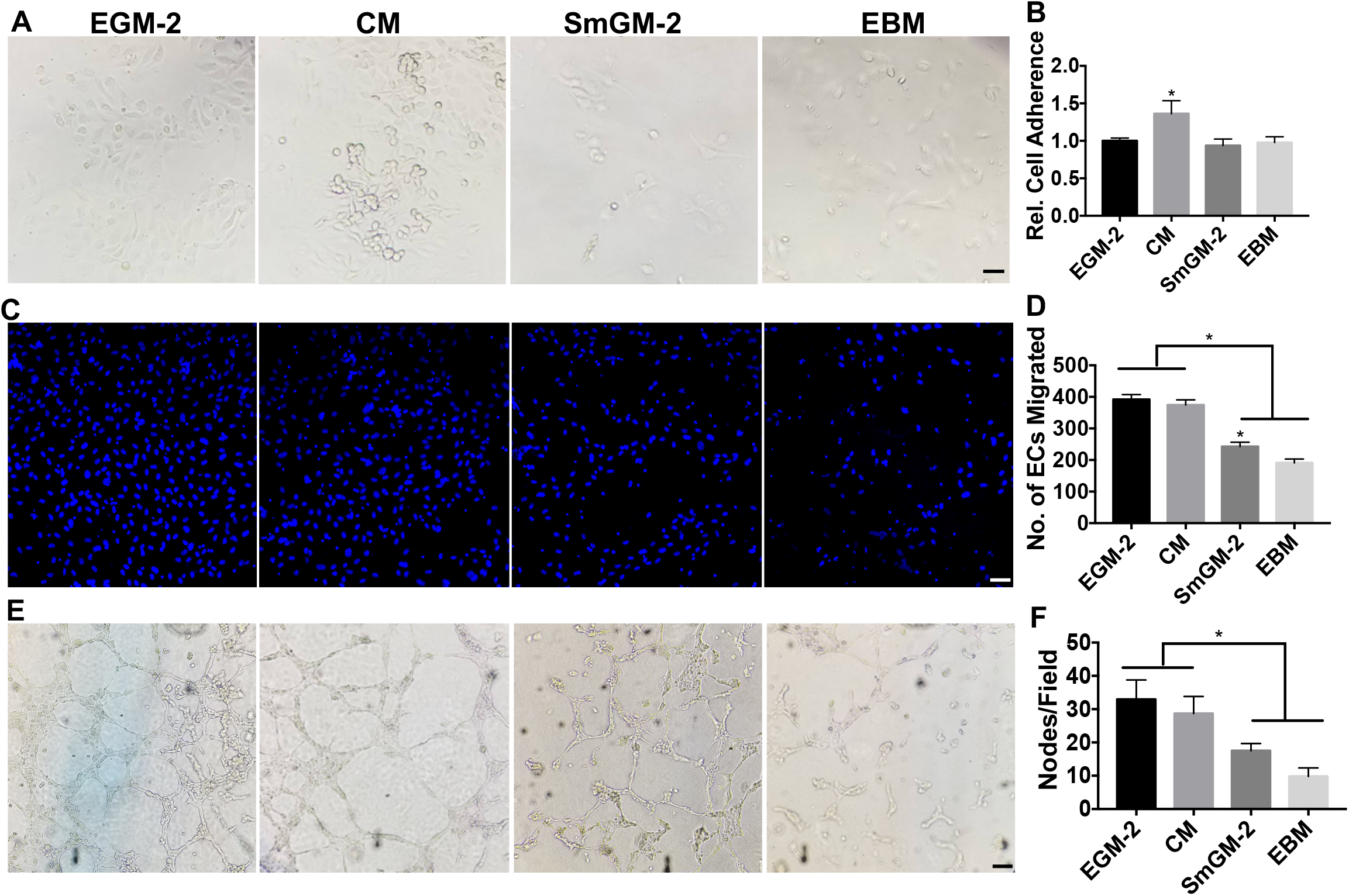
Functional assessment of the conditioned medium from collagen/fibronectin scaffolds. The conditioned medium (CM) collected from the collagen/fibronectin scaffolds were tested for their proangiogenic bioactivity using HUVEC. (A) Brightfield images showing attachment of HUVEC on 0.1% coated gelatin wells after 3 hr of seeding. Scale bar measures 100 μm. (B) AlamarBlue assay was performed to determine relative level of HUVEC adherence on gelatin coated plate after 3 hr of seeding. Eight micrometer pore size trans-wells were used to determine migration of HUVECs in response to the CM over 4 hr of incubation time. (C) The cells migrated were stained with dapi and (D) counted to obtain the total number of cells migrated. Scale bar measures 50 μm. Angiogenesis assay was performed using Matrigel and HUVEC. (E) Brightfield images showing network formation of HUVEC in response to CM from in situ collagen/fibronectin scaffolds after 6 h of incubation. Scale bar measures 100 μm. (F) The graph represents number of nodes/field. Endothelial cell (EGM-2) and smooth muscle cell (SmGM-2) growth medium, and endothelial cell basal medium (EBM) were used as controls. * denotes statistical significance differences between the different groups (n=3-6, one-way ANOVA, *p<0.05).

## Discussion

Human iPSC-derived VSMC has lately received major scientific attention in regenerative vascular research [7]. Recent attempts to generate robust implantable blood vessels and regenerate wounds have rendered these cells highly relevant for the development of functional vascular grafts and proangiogenic therapeutic modules [12, 16, 17, 27, 28]. An earlier study has implicated LM-VSMC in promoting angiogenesis [15]. We have taken a step forward and investigated the proangiogenic and immunomodulatory secretory function of this hiPSC-VSMC from LM embryonic origin for wound healing application [16, 17].

In our recent studies, we reported that the hiPSC-VSMC delivered in a collagen-based biomimetic scaffold promotes wound closure with enhanced vascular regeneration and reduced inflammation [16]. We attempted to understand how extracellular matrix such as collagen modulates their function. Our study led to the understanding of the effect of collagen fibrillar density on the paracrine secretion of hiPSC-VSMC. We showed that increase in collagen fibrillar density enhances the secretion of major proangiogenic factors such as vascular cell endothelial growth factor (VEGF), bFGF, angiopoietin (ANG)-1, platelet-derived growth factor (PDGF), transforming growth factor (TGF)-β1, stromal cell-derived factor (SDF)-1α, and MMP-2 [16]. In another study, we tried to understand the effect of hyaluronic acid (HA) on the hiPSC-VSMC secretory function. The HA while promoted proliferation of hiPSC-VSMCs in a collagen scaffold did not show any effect on their secretory function [25].

This sets the stage for developing strategies for precise modulation of hiPSC-VSMC’s function. One of the approaches is the development of integrin-targeting biomaterials to modulate the cell’s function for improved regenerative potential [19]. The method involves functionalization of the biomaterials with ECM biomolecules with a central idea that the ligand of the ECM biomolecules targets specific integrin interactions and regulate cellular responses, such as increased cell migration, proliferation, and paracrine secretion [22, 29, 30]. This hiPSC-VSMC from LM origin has earlier shown to be expressing integrin β3 and was found to be highly upregulated in proliferative phenotype [14]. In this study, we functionalized the collagen scaffold with a biomolecule that can facilitate specific interaction with the integrin β3 receptor. One of the natural choices was fibronectin. Fibronectin has two binding sites one is an integrin-binding site and the other is RGD or growth factor (GF) binding site next to it [20–22]. The simultaneous binding of GF and integrins can lead to a synergistic integrin/GF receptor signaling enhancing the effect of GF on hiPSC-VSMC.

Our initial screening on 2D tissue culture plates showed improvement in cell proliferation of hiPSC-VSMC in response to fibronectin. This was further validated using an antagonist of αvβ3, echistatin. In a 3D setup fibronectin in a fibrillar collagen scaffold enhances cell proliferation and secretion of bFGF. We were able to reverse the rate of proliferation and bFGF secretion by inhibiting αvβ3 integrin signaling. Fibronectin regulates the function of the cell through α5β1 as well. However, we did not see fibronectin regulating cell proliferation and bFGF secretion via α5β1 as shown by incubating with ATN-161, an antagonist of α5β1. Furthermore, the fibronectin regulates the cell proliferation and bFGF secretion in a collagen fibril density-dependent manner as shown by the 4mg/ml collagen density with enhanced cell proliferation and bFGF secretion. Most importantly, the amount of fibronectin determined the proliferation and bFGF secretion. The higher the amount of fibronectin the better the response while the lower amount of fibronectin such as 10 and 25 μg/ml did not elicit the desired response. The cells were shown to be proliferative as shown in their elongated morphology in the fibronectin functionalized scaffold.

FGFs and their role in angiogenesis are very well studied. bFGF is one of the FGFs which acts as a potent inducer of angiogenesis [31]. bFGF positioned at the upstream in the signaling system interacts with growth factors and chemokines such as VEGF, PDGF, hepatocyte growth factor, and monocyte chemoattractant protein to contribute towards the development of mature vessels and collateral arteries [32]. We believe an elevated level of bFGF in the case of the CM of the fibronectin functionalized scaffold might be responsible for promoting cell attachment, migration, and network formation of HUVECs in vitro. bFGF is also known to interact with αvβ3 integrin to promote vascular cell adhesion during angiogenesis [33]. We believe a similar mechanism might be existing here. bFGF and αvβ3 might be creating a positive feedback loop for enhanced iPSC-VSMC cell adhesion.

MMPs, extracellular proteinase, are targets that regulate cell behavior and have recently been used to modulate stem cell function [34]. MMPs not only degrade structural proteins of ECM but also cleave cell surface molecules such as integrins to change cell function [35, 36]. Numerous studies have shown cross-talk between MMP-2 and αvβ3 to promote stem cell adhesion, proliferation, and migration [37]. In blood vessels, they are found to be colocalized [38]. MMP-2 has also been shown to cleave fibronectin into small fragments to enhance the adhesion and migration of human melanoma cells by αvβ3 [39]. In this study, we found that in collagen scaffold higher than 1.25mg/ml the MMP-2 is upregulated in the presence of fibronectin. We believe that the increase in MMP-2 might be facilitating an augmented interaction between fibronectin and αvβ3 integrins present in iPSC-VSMCs [40]. Besides, MMP-2 is crucial in releasing bFGF from ECM to regulate cell migration [37, 41]. MMP-2 is also known to stimulate the interaction of VSMCs with newly formed ECM [42]. During pathophysiology’s such as hypertension, this interaction between VSMC and ECM triggers in phenotypic switching from a contractile phenotype to a synthetic one for the cells to migrate [42]. So in other words release of MMP- 2 might be regulating the migration of iPSC-VSMC via integrin signaling and disrupting growth factors bound to ECM.

However, the exact mechanism of MMP-2 regulation in the fibronectin functionalized 3D collagen scaffold is not known. In the past, it has been shown that MMP-2 gene expression could be induced via activator protein-1 after integrin-linked kinase activation after cell integrin and fibronectin engagement [43]. However, our study with the integrin antagonists did not show any changes in the amount MMP-2 indicating any changes in the MMP-2 level is independent of integrin and fibronectin interaction. An earlier study shows a direct regulation of MMP-2 by bFGF in endothelial cells [44]. In another study on prostate cancer cells associated with prostate fibroblasts or osteoblasts showed enhanced MMPs in the presence of bFGF and TGFβ1 [45]. Nevertheless, the integrin inhibition assay which resulted in lower level bFGF was not enough to show any direct link between bFGF and MMP- 2. To completely block the expression of bFGF to determine the role of bFGF in MMP-2 secretion we adopted an engineering approach and cross-linked the fibronectin and collagen using a 4S-StarPEG nontoxic cross-linker that binds to free primary amine groups. Our idea was to make the binding motifs of fibronectin less accessible to integrin present in iPSC-VSMCs. The cross-linked resulted in a huge drop in the release of bFGF. We also saw a subsequent decrease in MMP-2 expression. This demonstrates that MMP-2 expression in hiPSC-VSMC is regulated by bFGF.

Finally, it is the combination of physical cues, the production of growth factors, and the MMPs that support hiPSC-VSMC proliferation in a fibronectin functionalized scaffold. Our study indicates the presence of a positive feedback loop between integrin β3, bFGF, and MMP-2. We, therefore, propose an IGM cycle (Integrin- Growth factor-MMP) to account for the modulation of hiPSC-VSMC (Figure 8).

**Figure 8:**
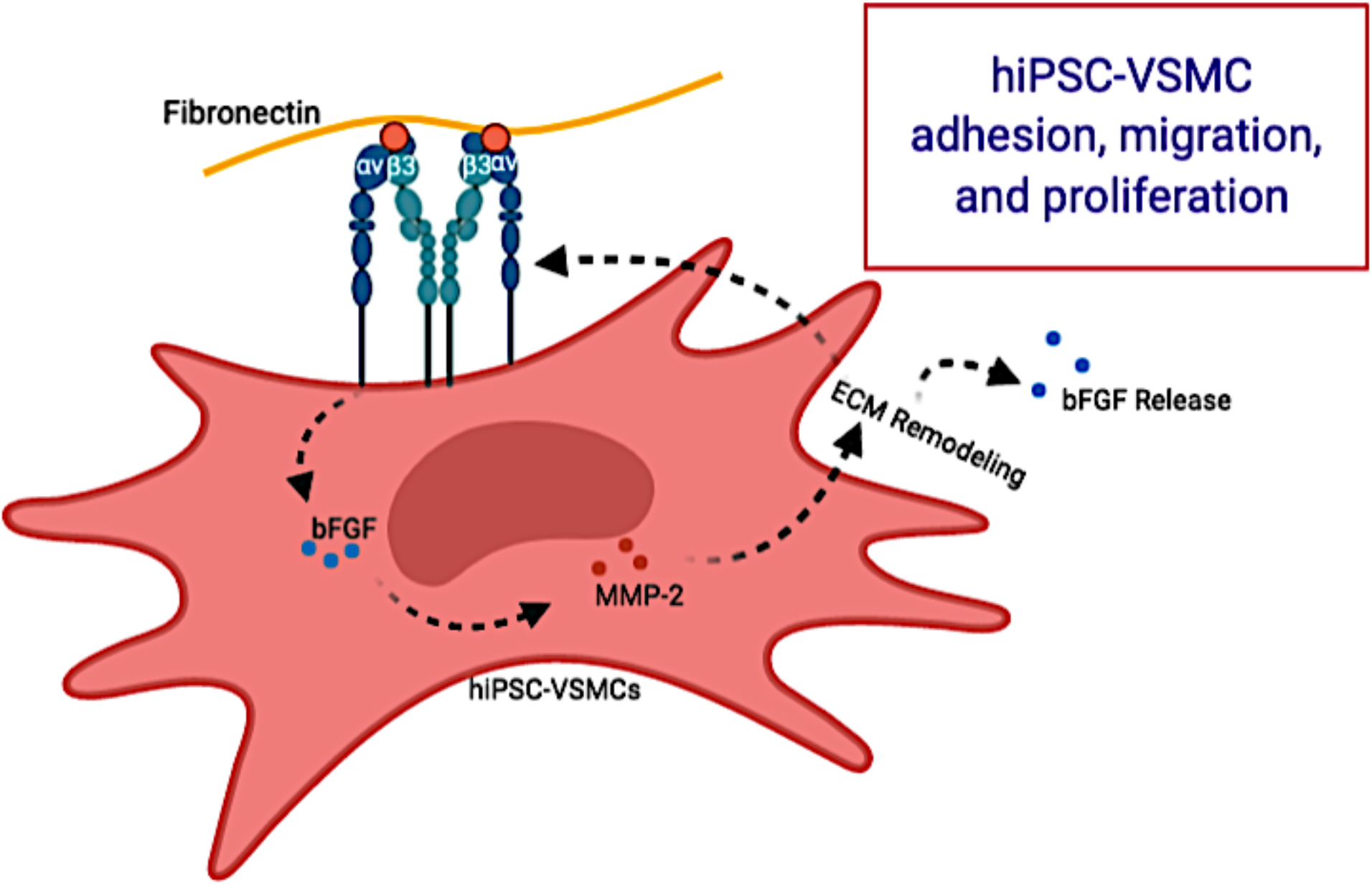
Schematic showing integrin β_3_, bFGF and MMP-2 positive feedback loop system modulating hiPSC-VSMC function. Created with BioRender.com

## Conclusion

In summary, we showed an approach to regulate the secretory function of hiPSC-VSMC by targeting their integrin β3 receptor. The fibronectin within a collagen scaffold promoted bFGF secretion via αvβ3 secretion and was dependent on the collagen fibrillar density and amount of fibronectin. Finally, we believe i) bFGF regulates the upregulation of MMP-2 in hiPSC-VSMCs in an integrin independent pathway and ii) a positive feedback loop between integrin β3, bFGF, and MMP-2 exists and promotes iPSC-VSMC proliferation. Furthermore, bFGF in the CM showed proangiogenic potential by enhancing cell adherence, migration, and network formation of HUVECs. Ongoing efforts are to engineer complex ECM-based biomaterial systems for developing iPSC-based vasculature.

## Supplementary Information

The Supporting Information is available online and contains additional information on materials, antibodies, and TNBSA method and data.

## Author Contributions

B.C.D. conceived the study. B.C.D. and H.C.H. procured the funding. B.C.D. designed the experiments. B.C.D. and K.D. performed the experiments. B.C.D. wrote the manuscript. All the authors participated in data analysis, discussed the results, and reviewed the manuscript.

## Funding

This work was funded by the Plastic Surgery Foundation Grant 18-003032 (H.C.H and B.C.D) and Yale Department of Surgery (H.C.H).

## Acknowledgment

The authors would like to thank the core research facilities at the Yale Department of Surgery.

## Conflicts of Interest

The authors declare no conflict of interest.

## Notes

### Competing Interest Statement

The authors have declared no competing interest.

