## Supplemental Information for "Integrin β3 Targeting Biomaterial Preferentially Promotes Secretion of bFGF and Proliferation of iPSC-Derived Vascular Smooth Muscle Cells"

**Materials:**

Type I rat tail collagen type-I was purchased from Enzo Life Sciences, USA. Human plasma fibronectin was purchased from Milipore Sigma, USA. 4-arm polyethylene glycol succinimidyl glutarate Mw 10,000 (4S-StarPEG) was purchased from JenKem Technology, USA. Trinitrobenzene sulfonic acid (TNBSA) and AlamarBlue reagents were purchased from ThermoFisher Scientific, USA. Smooth muscle cells culture medium (SmGM-2) was purchased from PromoCell, Germany. All other cell culture reagents were purchased from ThermoFisher Scientific unless otherwise stated.

**2,4,6-trinitrobenzene sulfonic acid assay**:

2,4,6-Trinitrobenzene Sulfonic Acid (TNBSA) assay was performed to determine the amount of free amines in the scaffolds with and without cross-linking. Briefly, the scaffolds were first made inside Eppendorf tubes following the supplementary table described above. Then the scaffolds were disrupted using crushers, which would turn the scaffold inside a viscous consistency. 250uL of 0.01% TNBSA was added into 500uL of each sample. A 0.2% glutaraldehyde solution was used as a positive control. The resultant mixtures were incubated at 37^o^C for 2 hours. Glycine titration in sodium bicarbonate (0.1M) pH 8.5 standard curve was established at: 300 µm, 250 µm, 200 µm, 150 µm, 100 µm, 75 µm, 50 µM, 25 µm, and 0 µm. At the conclusion of the incubation, 250 µl of 10% sodium dodecyl sulfate was added. 125 µl of 1M HCI was added to stop the reaction. 100 µl of the final solution was used to test for the degree of free amine groups in fibronectin infused collagen scaffolds. Plate reader absorbance was set at 335 nm.

**Table-S1:** Information related to primary antibodies.

| Primary Antibody | Dilution | Catalog Number/ Manufacturer |
| --- | --- | --- |
| SDF-1α | 1:2500 ELISA | MAB350 (R&D) |
| PDGFAA | 1:2500 ELISA | 500-P46-100 (PeproTech) |
| bFGF | 1:2500 ELISA | 500M38 (PeproTech) |
| VEGF | 1:2500 ELISA; 1:200 IHC | AB-119 (Abcam) |
| SM-22α | 1:300 IF and IHC  1µg/ml FACS | AB-10135 (Abcam) |
| Calponin | 1:200 IF and IHC  1µg/ml FACS | C-2687 (Sigma) |

| Secondary Antibody | Dilution | Catalog Number/ Manufacturer |
| --- | --- | --- |
| Anti-Mouse-HRP | 1:2500 ELISA | AB6789 (Abcam) |
| Anti-Rabbit-HRP | 1:2500 ELISA | A0545 (Sigma) |
| Anti-Mouse-Alexafluor488 | 1:400 IF and IHC | A11029 (ThermoFisher) |
| Anti-Goat-Alexafluor488 | 1:400 IF and IHC | A27012 (ThermoFisher) |

**Table-S2:** Information related to secondary antibodies.

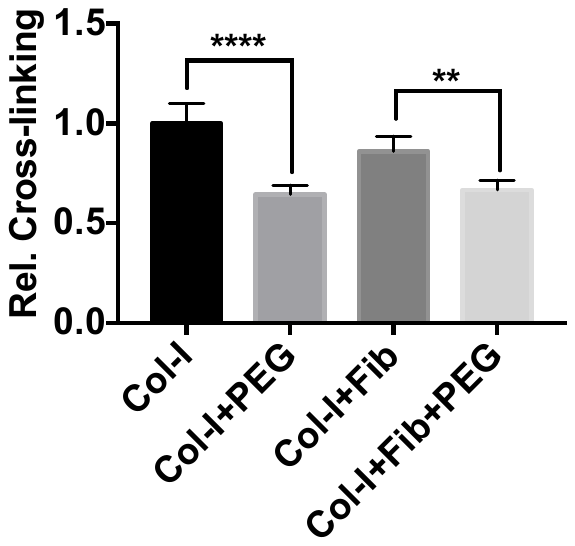

**Figure S1: Characterization of 4S-StarPEG cross-linking.** TNBSA showing a qualitative analysis of free amine groups in crosslinked collagen sacffolds of 4mg/ml of collagen concentration with and without fibronectin (100 µg/ml). Uncross-linked collagen scaffolds with and without fibronectin were kept as a controls. * denotes statistical significance differences between the different groups (n=4, t-test, **p<0.01, ****p <0.0001).
